# Coinfection with Intestinal Parasite Expands Resident Macrophages and Impairs Control of Chronic Herpesvirus Infection

**DOI:** 10.1101/2022.10.05.510926

**Authors:** Christina M. Zarek, Chaitanya Dende, Jaime Coronado, Mihir Pendse, Phillip Dryden, Lora V. Hooper, Tiffany A. Reese

**Affiliations:** Department of Immunology, University of Texas Southwestern Medical Center, Dallas, TX; Department of Microbiology, University of Texas Southwestern Medical Center, Dallas, TX

## Abstract

In addition to a range of homeostatic functions, resident macrophages are essential for immune surveillance in tissues. Therefore, anything that alters the phenotype or function of these cells potentially impacts their response to infectious challenges. Parasite infections cause proliferation of large peritoneal macrophages (LPMs), which are the resident macrophages of the peritoneal cavity. However, the functional consequences of LPM expansion on the control of secondary infectious challenge is unknown. Using a coinfection model with the intestinal parasite *Heligmosomoides polygyrus* (HP) and the virus, murine gammaherpesvirus-68 (MHV68), we investigated the impact of LPM expansion on viral infection. We determined that LPM expansion induced by HP required retinoic acid signaling. When we challenged HP-infected mice with MHV68, we observed increased herpesvirus infection and latency. Coinfection of mice with macrophage-specific deletion of GATA6, the retinoic acid-responsive transcription factor that drives LPM transcriptional programming, eradicated the increase in viral infection. In addition to increased MHV68 infection, parasite coinfected mice displayed increased herpesvirus reactivation from latency, indicating impaired control of chronic herpesvirus infection. Elimination of dietary vitamin A, which depletes retinoic acid and LPMs, abolished the increased MHV68 reactivation in parasite coinfected mice. These results indicate that parasite- and retinoic acid-mediated resident macrophage expansion drives increased herpesvirus infection, latency, and reactivation.

## Introduction

Helminth infections are endemic in many parts of the world, and parasites have systemic immunomodulatory roles. Parasites, such as the mouse helminth, *Heligmosomoides polygyrus* (HP), reside exclusively in the small intestines of infected animals, but modulate the immune responses to other pathogens in non-intestinal tissues (*1*–*3*). They do this, in part, by expanding regulatory T cell populations, skewing cytokine profiles, and excreting immunomodulatory products. Adaptations like these allow helminths to survive in their hosts for extended periods, but how these adaptions affect the immune system’s ability to respond to a secondary pathogen is less well studied.

One intriguing feature of parasite infections is their ability to expand tissue resident macrophages, particularly in the serous cavities. In the peritoneal cavity, there are two main populations of macrophages: large peritoneal macrophages (LPMs) and small peritoneal macrophages (SPMs). LPMs are self-renewing, fetal progenitor-derived tissue resident macrophages and are important for immune homeostasis, such as clearing apoptotic cells (*4*–*6*). Various species of helminth, including *Heligmosomoides polygyrus* (HP), cause proliferation of the LPM population in an IL-4 and CSF1R-dependent manner (*7*–*9*). The impact of this proliferation has not been elucidated, but one possible outcome may by increased control of helminth infection (*10, 11*). In addition, this expansion of tissue resident macrophages in various tissues of the body during intestinal helminth infection represents a potential novel mechanism of immunomodulation.

Gammaherpesviruses, like many helminths, are chronic pathogens. Murine gammaherpesvirus-68 (MHV68) is a natural rodent pathogen that models gammaherpesvirus pathogenesis in a mouse because of its close genetic relationship with the human herpesviruses Epstein Barr virus and Kaposi’s sarcoma-associated herpesvirus. After acute replication, MHV68 establishes latency in B cells, macrophages and dendritic cells and persists for the remainder of the host’s life. Certain signals, such as HDAC inhibitors (*12*), parasite infection (*13*), or IL-4 complexed with IFN-γ blockade (*14*), promote virus reactivation from latency, which is important for herpesvirus-induced lymphomas (*15*).

Interestingly, MHV68 was recently shown to infect LPMs (*16*), leading us to question what the impact of parasite-mediated expansion of LPMs would be on peritoneal viral infection. Studies examining the outcomes of parasite and gammaherpesvirus coinfection reveal the complex nature of coinfection studies. We showed previously that if mice infected with a chronic MHV68 infection are challenged with *H. polygyrus* or *Schistosoma mansoni* eggs, the virus reactivates from latency in an IL-4-dependent manner specifically from macrophages (*13, 14*). However, when mice are first treated with *Schistosoma mansoni* eggs, then challenged with MHV68 intranasally 14-days later, the virus replicates less in the lung (*17*). Species of helminth, timing of infection, immune response to the pathogens, and tissue tropism all play a role in whether the coinfection helps or hinders the host response to the virus (*18*).

Recognizing that the timing of coinfections and the tissue examined are critical variables, we established a system of intestinal parasite infection with *Heligmosomoides polygyrus*, followed by intraperitoneal challenge with MHV68. Using this system, we sought to gain a better understanding of the regulation of LPMs during intestinal helminth infection and to elucidate the impact of expanded LPMs during coinfection with a virus.

## Results

### LPMs expand during *H. polygyrus* infection independently of STAT6, but dependent on retinoic acid

To investigate the regulation of LPMs after intestinal helminth infection, we challenged mice orally with 100 L3 larvae of *H. polygyrus* (HP). When we challenge mice with HP, we observed expansion of LPMs, similar to previous reports. We found that HP-infected mice have increased numbers of LPMs at days 7 and 14 post infection. LPM numbers returned to uninfected levels by day 34 post infection (Figure 1A, Figure 1 supplement). Previous reports of helminth-induced tissue resident macrophage expansion demonstrated that signaling through the IL-4 receptor was sufficient to cause expansion of LPMs (*7, 8*). Moreover, delivery of long-lasting IL-4 complexes induced LPM expansion (*8, 19*). To determine if HP-induced LPM expansion is dependent on IL-4 receptor signaling, we infected *Stat6*^*-/-*^ or littermate control mice with HP and measured the number of LPMs. In contrast to the previous studies with the parasites *Litomosoides sigmodontis* and *Schistosoma mansoni*, we found that HP-induced LPM expansion did not depend on IL-4 signaling through STAT6, as *Stat6*^*-/-*^ mice still had normal expansion of LPMs (Fig 1B). This suggests that LPMs require signals other than IL-4 or IL-13 to induce their expansion in response to HP.

**Figure 1.**
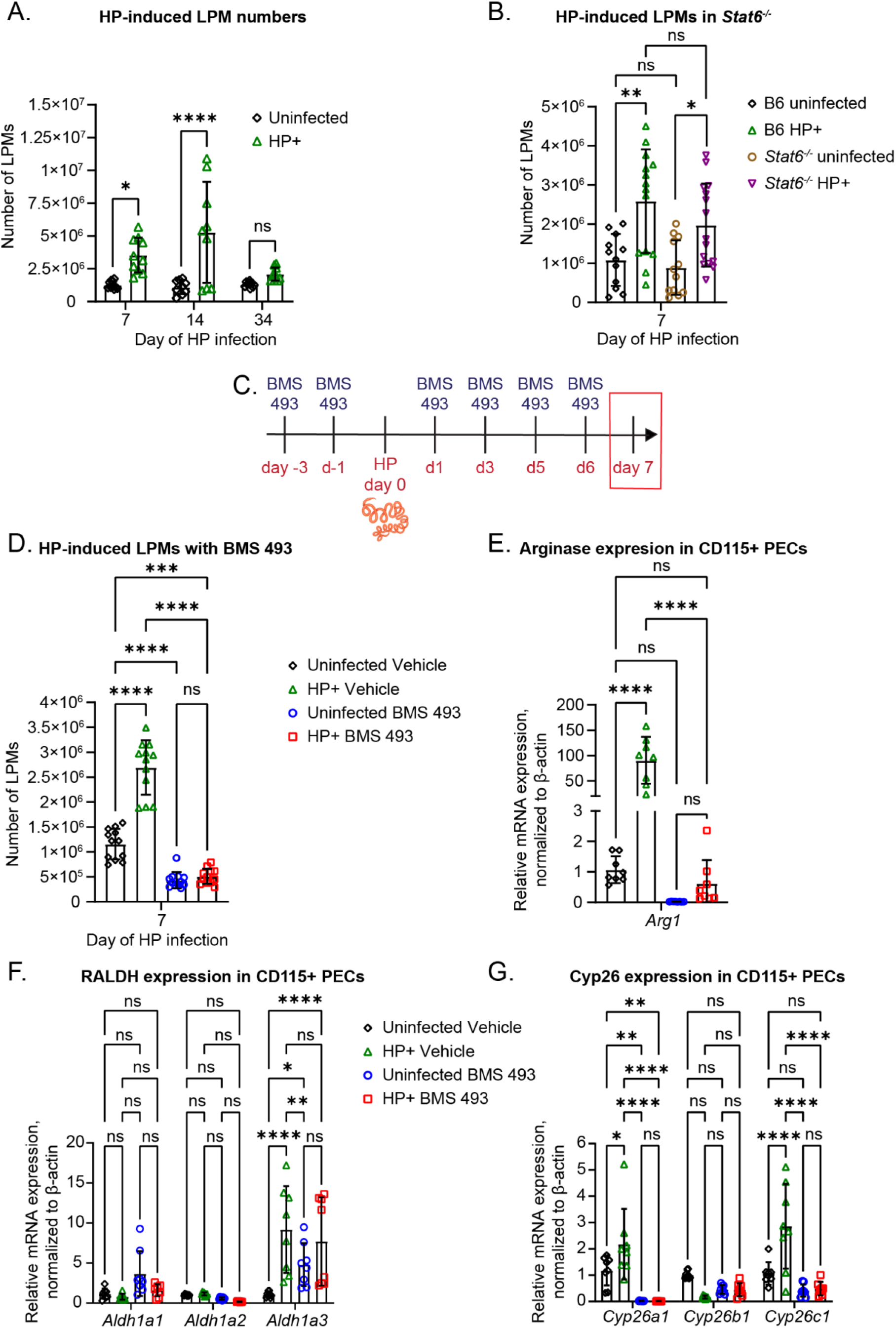
LPMs expand during *H. polygyrus* infection independently of STAT6, but dependent on retinoic acid. C57BL/6J mice were infected with 100 L3 larvae of HP by oral gavage. (A) Quantification of flow cytometric analysis of LPMs in C57BL/6 mice at the indicated days post HP infection. LPMs were gated as CD3-CD19-Siglec F-CD11b^hi^ ICAM-2^hi^. Data are pooled from 3 independent experiments (n=9-10/group, mean ± standard deviation). (B) Quantification of flow cytometric analysis of LPMs in C57BL/6 and *Stat6*^-/-^ mice on day 7 of HP infection. Data are pooled from 3 independent experiments (n=12-15/group, mean ± standard deviation). Each dot represents an individual mouse. (C) Timeline of BMS 493 oral gavage treatments and HP infection. Partially created in BioRender. (D) Quantification of flow cytometric analysis of LPMs in C57BL/6 mice treated with BMS 493 or vehicle seven days after HP infection. LPMs were gated as CD45+ CD3-CD19-Siglec F-CD11b^hi^ ICAM-2^hi^. Data are pooled from 3 independent experiments (n=12/group, mean ± standard deviation). (E-G) CD115+ PECs were purified with MACS columns on day 7 after HP infection before RNA extraction. CT values were normalized to β-actin and then normalized to the delta CT of uninfected vehicle treated mice. Data pooled from 2 independent experiments. (n=8 mean ± standard deviation). (E) CD115+ PEC expression of a RA-dependent gene, *Arg1*. (F) Expression of RALDH genes in CD115+ PECs. (G) Expression of Cyp26 genes in CD115+ PECs. P-values, 2-way ANOVA, Tukey’s multiple comparisons. * P ≤ 0.05, ** P ≤ 0.01, *** P ≤ 0.001, **** P ≤ 0.0001

Because retinoic acid, a derivative of dietary vitamin A, is important in LPM localization, polarization, and proliferation (*20, 21*), we next asked whether retinoic acid responsive genes were changed by HP infection. We looked at expression of a retinoic acid dependent gene, *Arg1* (*20, 21*). We also examined genes that are induced by retinoic acid signaling and are important for the metabolism of all trans-retinoic acid (ATRA), the *Aldh1a* and *Cyp26* genes. *Aldh1a* genes code for retinaldehyde dehydrogenases (RALDH), the enzymes that convert retinaldehyde to ATRA, while cytochrome P450 (Cyp26) enzymes convert ATRA to less biologically active forms. We isolated CD115+ PECs seven days after HP infection. In HP-infected mice we observed an increase in expression of *Arg1, Aldh1a3, Cyp26a1* and *Cyp26c1* in the CD115+ PECs (Fig. 1E, F, G), which suggests that genes that are induced by retinoic acid were increased in macrophages in the peritoneal cavity after intestinal parasite infection.

We next asked whether retinoic acid was required for LPM expansion after HP infection. To address this, we used a pan retinoic acid receptor (RAR) inverse agonist, BMS 493 (*22, 23*). RARs are nuclear receptors that when bound to retinoic acid, regulate the expression of retinoic acid target genes. Blockade of RARs with BMS 493 leads to increased association of RARs with corepressors and decreased recruitment of transcriptional coactivators to retinoic acid responsive genes. Mice were treated with BMS 493 or vehicle control starting 3 days prior to HP infection and continuing every other day during the course of infection (Fig. 1C). We measured LPMs after BMS 493 treatment and observed decreased numbers of LPMs compared with vehicle treated mice (Fig. 1D). Further, LPMs did not expand in mice that were infected for seven days with HP and treated with BMS 493 (Fig. 1D). When we quantitated gene expression in macrophages from the BMS 493-treated mice, we observed that BMS 493 countered the induction of retinoic acid-induced genes, *Cyp26a1, Cyp26c1*, and *Arg1* in HP-infected mice (Fig. 1E, F, G). We noted that *Aldh1a3* was induced by BMS 493 relative to vehicle treated mice, and that HP infection did not further induce *Aldh1a3* in BMS 493 treated mice. Although *Aldh1a3* is retinoic acid-responsive it is also possible that deprivation of retinoic acid may induce expression of RALDH enzymes because they encode for the rate limiting enzymes that produce ATRA. Taken together, these data indicate that BMS 493 treatment, which antagonizes retinoic acid induced gene expression and metabolism, abolished LPM expansion after HP infection, suggesting that LPM expansion after intestinal parasite infection was regulated by retinoic acid.

Our data combined with the data from others establish that many helminths induce tissue resident macrophage expansion. However, the consequences of this expansion on secondary infections are still unclear. In particular, the impact of LPM expansion on peritoneal viral infection has never been reported, even though we routinely model viral infection in mice by peritoneal injection. Murine gammaherpsvirus-68 (MHV68) infects and establishes latent, chronic infection in B cells, dendritic cells, macrophages, and in particular, large peritoneal macrophages(*16, 24, 25*). We asked whether HP infection altered acute infection with MHV68. To establish coinfection, we first infected with 100 L3 HP larva by oral gavage, then intraperitoneally injected 1 × 10^6^ PFU of MHV68 one week later (Figure 2A).

**Figure 2.**
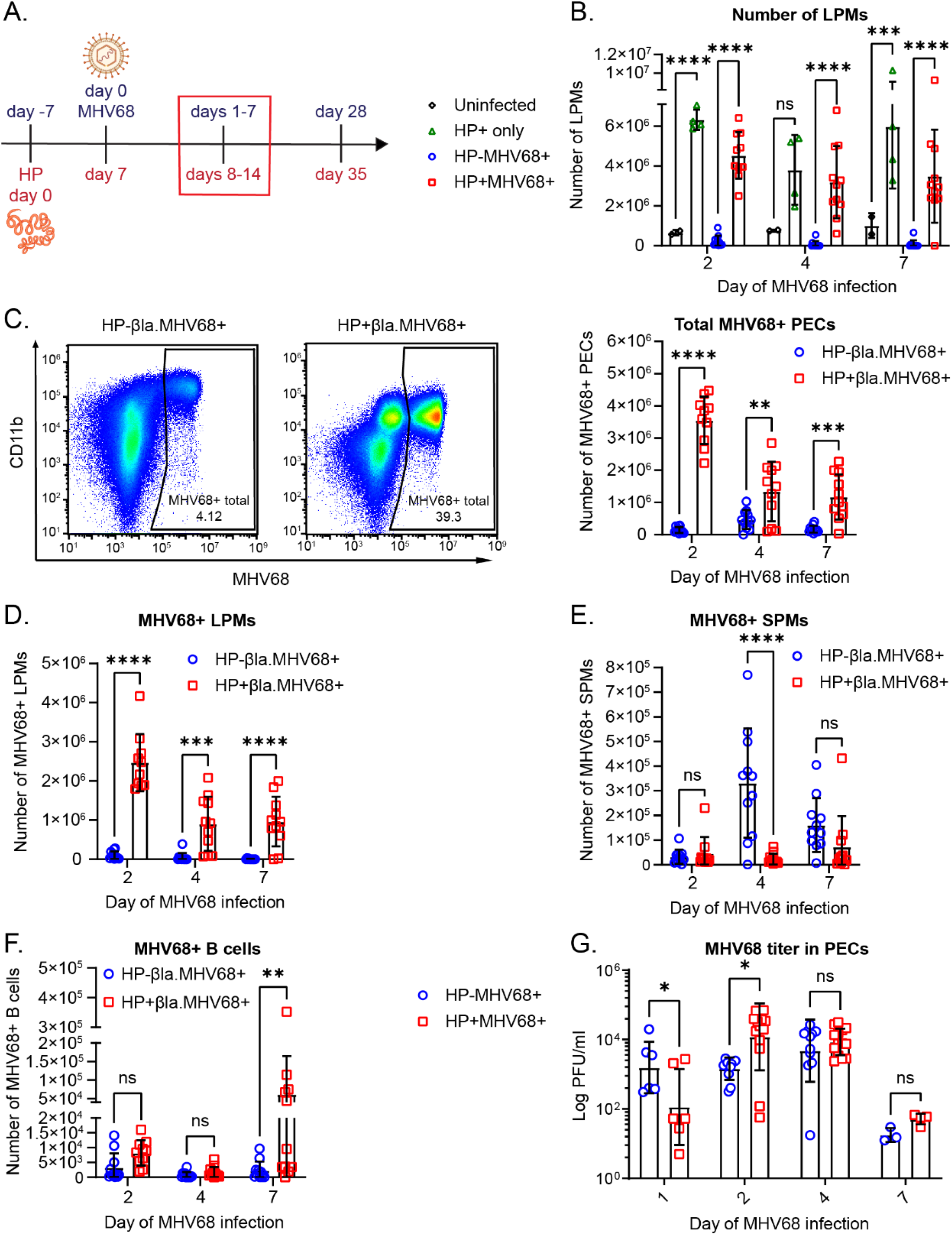
LPM expansion during intestinal parasite infection increases viral infection in the peritoneal cavity. (A) Timeline of infections with HP by oral gavage and MHV68 by intraperitoneal injection (10^6^ plaque forming units (PFU)). The time points shown in (B-G) are outlined by a red box. Partially created in BioRender. (B) Quantification of flow cytometric analysis of LPMs in C57BL/6 mice at days 2, 4, and 7 post MHV68 infection. LPMs were gated as CD19-CD11b^hi^ ICAM-2^hi^. Data are pooled from 3 independent experiments (n=10-11 MHV68+ groups, n=4 HP+ only group, n=2 uninfected group, mean ± standard deviation). (C-F) Mice were infected as in A with 10^6^ PFU of MHV68.ORF73β-lactamase virus. Quantification of flow cytometric analysis of MHV68-infected PECs at days 2, 4, and 7 post MHV68 infection. Data are pooled from 3 independent experiments (n=10-11/group, mean ± standard deviation). (C) Representative flow plots of the total β-lactamase-positive (MHV68+) PECs. Quantification of the total number of MHV68-infected PECs. (D) Number of MHV68-infected LPMs. (E) Number of MHV68-infected SPMs. SPMs were gated as CD19-CD11b+ ICAM-2^lo^. (F) Number of MHV68-infected B cells. B cells were gated as CD19+. Each dot represents an individual mouse. (G) MHV68 titers determined by plaque assay of PECs at day 1-7 of infection. Data are pooled from 5 independent experiments (n=6-7 day 1, n=11-14 days 2 and 4, n=3-4 day 7, mean ± standard deviation). P-values, 2-way ANOVA, Tukey’s multiple comparisons * P ≤ 0.05, ** P ≤ 0.01, *** P ≤ 0.001, **** P ≤ 0.0001

This timeline allows the HP to establish an active infection before introducing the virus and ensures that LPMs are expanded at the time of MHV68 infection. We first assessed LPM expansion over a time course in HP/MHV68 coinfected mice compared with uninfected and single infected mice. We collected PECs at day 2, 4, and 7 of MHV68 infection and performed flow cytometry. We found that coinfected mice had expansion of LPMs, comparable to HP-only infected mice. In contrast, mice infected with MHV68 only had decreased numbers of LPMs (Figure 2B).

Since we wished to examine specifically MHV68 infection in LPMs and other cells, we infected mice with a reporter virus, MHV68.ORF73β-lactamase. This virus encodes β-lactamase fused to mLANA (ORF73), which is active during both acute replication and latency (*26*). This means that both cells with replicating virus and latent virus will be marked by the reporter virus. We found that the total number of MHV68-infected PECs was increased in coinfected mice compared to mice infected only with virus (Figure 2C). Further, there were increased numbers of MHV68-infected LPMs in coinfected mice (Figure 2D). We also measured MHV68 infection in the monocyte-derived SPMs and noted that the number of MHV68-infected SPMs tended to be higher in virus-only infected mice at days 4 and 7 of infection compared to the coinfected mice (Figure 2E). When we examined infected B cells, we noted that HP coinfection also led to increased B cell infection at day 7. However, there were more than 10-fold fewer infected B cells than macrophages in HP/MHV68 coinfected and MHV68-single infected mice, suggesting that the majority of infected cells were LPMs (Figure 2F). These data indicate that MHV68 infection at early timepoints occurred primarily in LPMs, in agreement with previous reports (*16*). In addition, parasite coinfection significantly expanded the number of infected LPMs during acute viral infection.

Since we saw increased MHV68 infection in coinfected mice, we questioned whether there was increased replication of MHV68, as well. To test this, we collected PECs and performed plaque assays at days 1, 2, 4, and 7 of MHV68 infection (Figure 2G). We found that coinfected mice had decreased titer at day 1 of infection, but increased titer at day 2 of infection. By day 4, the titers between the virus-only infected mice and coinfected mice were not significantly different, and this was maintained at day 7 of infection. When compared to the infection data with the marker virus, this suggests that even though there was increased MHV68-infected cells in the coinfected mice, that these infected cells were not replicating more virus and instead harbor latent virus. This observation supports the idea that some MHV68 genomes become latent immediately (*16, 27*). Overall, our data shows that coinfected mice have increased numbers of infected LPMs.

### Coinfected mice have increased MHV68 infection in tissue resident macrophages during latency

Since we saw that coinfected mice had increased numbers of MHV68-infected PECs during acute replication, we questioned if this was maintained during latency. To address this, we infected mice and collected PECs for flow cytometry at day 28 of MHV68 infection, when the virus is latent and there is no remaining acute replication (Figure 2A). First, we examined the macrophage composition of the peritoneal cavity. We found that the LPM population was increased in the coinfected mice compared to MHV68-only infected mice (Figure 3A). The number of SPMs was equivalent between the MHV68-only and HP/MHV68 coinfected mice (Figure 3A). We confirmed that herpesvirus infection did not impact helminth infection, by quantitating worm burden and fecundity in both helminth-only infected and coinfected mice. We found that there was no difference in worm burden or fecundity when mice were coinfected with virus (Supplementary Figure 2A, 2B). These data suggest that helminth coinfection led to prolonged expansion of the LPM population in the peritoneal cavity of MHV68 infected mice.

**Figure 3.**
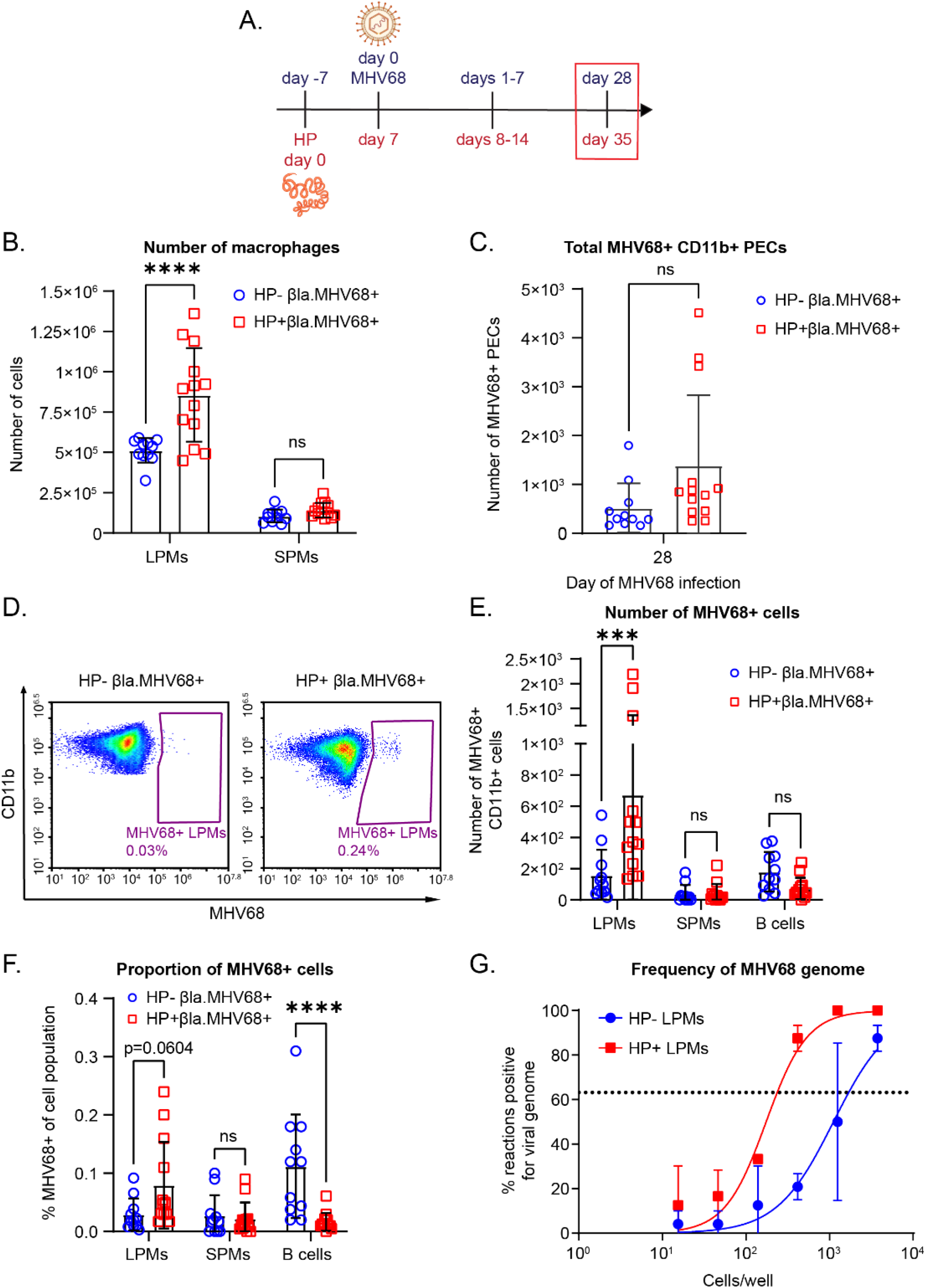
Coinfected mice have increased infection in tissue-resident macrophages during latency. (A) Timeline of infections with HP by oral gavage and MHV68 by intraperitoneal injection. The timepoints shown in (B-F) are outlined by a red box. Partially created in BioRender. (B) Quantification of flow cytometric analysis of the number of LPMs and SPMs at day 28 of MHV68 infection. Data are pooled from 3 independent experiments (n=11-13/group, mean ± standard deviation). Each dot represents an individual mouse. (C-F) Quantification of flow cytometric analysis of MHV68-infected CD11b+ PECs at day 28 of MHV68 infection. CD11b+ cells were isolated with CD11b+ beads and MACs columns before staining. LPMs were gated as CD19-CD11b^hi^ ICAM-2^hi^. SPMs were gated as CD19-CD11b+ ICAM-2^lo^. B cells were gated as CD19+. Data are pooled from 3 independent experiments (n=11-13/group, mean ± standard deviation). Each dot represents an individual mouse. (C) Total number of MHV68-infected CD11b+ PECs. (D) Representative flow plots of MHV68+ LPMs. (E) Quantification of number of MHV68-infected macrophages and B cells. (F) Proportion of MHV68-infected cells out of the parent populations. (G) LPMs were sorted from mice at days 28 and 29 of MHV68 infection as CD19-SiglecF-Ly6G-F4/80+ CD11b^hi^ ICAM-2^hi^. Sorted cells were subjected to limiting dilution PCR analysis to detect frequency of viral genomes. Data are pooled from 2 independent experiments (4-8 mice were pooled for each group before sorting, mean± standard deviation). (B, D, E) P-values, 2-way ANOVA, Tukey’s multiple comparisons. (C) P-values, unpaired t-test. (G) Dotted line represents Poisson distribution. * P ≤ 0.05, ** P ≤ 0.01, *** P ≤ 0.001, **** P ≤ 0.0001.

Using the MHV68.ORF73β-lactamase reporter virus, we next determined the number of infected cells at day 28 of MHV68 infection. CD11b+ cells were isolated by MACs columns before staining for flow cytometry to enhance the ability to detect MHV68 infected cells. We found that the total number of MHV68-infected CD11b+ PECs was not significantly increased in coinfected mice during latency (Figure 3C). In contrast, when we examined MHV68 infection specifically in the LPMs, we found that the number of MHV68-infected LPMs was increased in coinfected mice compared to mice that received only virus (Figure 3D, 3E). The total number of infected SPMs and CD11b+CD19+ B1 B cells was equivalent between the two groups (Figure 3E), suggesting that helminth infection only induced increased infection in LPMs. We also found that the proportion of infected LPMs was higher in coinfected mice than mice that received only virus, indicating that the increased number of infected LPMs was not simply due to an increase in the number of available cells. There was no difference in the proportion of infected SPMs between the groups, but there was a decrease in the proportion of infected B1 B cells in coinfected mice (Figure 3F).

Because we saw increased infection at both acute and latent timepoints of MHV68 in HP-coinfected mice, we questioned whether LPMs have an increased frequency of latency in coinfected mice. To address whether LPMs from coinfected mice harbor more latent virus, we sorted LPMs from coinfected and singly infected mice and used those cells in limiting dilution PCR (LD-PCR)(*28*). LD-PCR measures the frequency of MHV68 viral genomes by nested PCR for a MHV68 gene, ORF72, and is sensitive down to one copy of viral genome. When we compared the point at which the dilution curves crossed Poisson distribution, we observed that the frequency of viral genome in sorted LPMs from coinfected mice was approximately 1 out of 237 LPMs, while the frequency of viral genome from mice infected only with virus was approximately 1 out of 1,698 LPMs (Figure 3G). This confirms that LPMs in coinfected mice have an increased frequency of MHV68 genome than LPMs from mice infected only with MHV68. These data demonstrate that coinfection with a helminth prolonged the expansion of the LPM population, and increased MHV68 infection and latency in the LPMs.

### HP-induced increased latent MHV68 infection is dependent on vitamin A

Given that our data indicated that retinoic acid signaling was required for LPM expansion after HP infection and others have shown that LPMs are dependent on vitamin A, the precursor to retinoic acid (*20*) we questioned whether the increased MHV68 infection of LPMs in coinfected mice was dependent on vitamin A. To address this, we utilized a dietary depletion of vitamin A. In this system, dams are placed on a vitamin A deficient (VAD) diet or control diet one week before pups are born. The pups are maintained on their respective diets until euthanasia. In accordance with previous work (*19, 20*), the mice on the VAD diet had severely reduced numbers of LPMs compared to mice on the control diet, but normal numbers of SPMs (Figure 4A, 4B). Worm burden and fecundity were not affected by the VAD diet, suggesting that loss of vitamin A does not affect HP infection (Supplementary Figure 3A, 3B). We infected the mice with MHV68.ORF73β-lactamase to determine if coinfected mice on the VAD diet had increased infection. At day 28 or 29 of MHV68 infection, we found that virus-only infected mice on both control diet and the VAD diet had similar numbers of total MHV68+ CD11b+ PECs, suggesting that vitamin A deficiency does not impact baseline MHV68 infection (Figure 4C). On the other hand, when we quantitated the total number of MHV68-infected CD11b+ cells in HP-coinfected mice on the VAD diet, we found no increase in virus infection compared to virus-only mice on the VAD diet. This suggests that the helminth-induced increased infection was dependent on vitamin A (Figure 4C). We further subdivided the infected CD11b+ cells into LPMs and SPMs. As expected, HP-coinfected mice fed the control diet had increased numbers and proportion of MHV68-infected LPMs (Figure 4D and 4E). In contrast, HP-coinfected mice on the VAD diet had no increase in virus-infected LPMs (Figure 4D and 4E). Further, there was no difference in the number of infected SPMs between coinfected and virus-only mice on either diet (Figure 4D and 4E). Overall, these results suggest that the increased number of MHV68-infected cells in HP-coinfected mice was dependent on vitamin A and the expansion of the LPMs.

**Figure 4.**
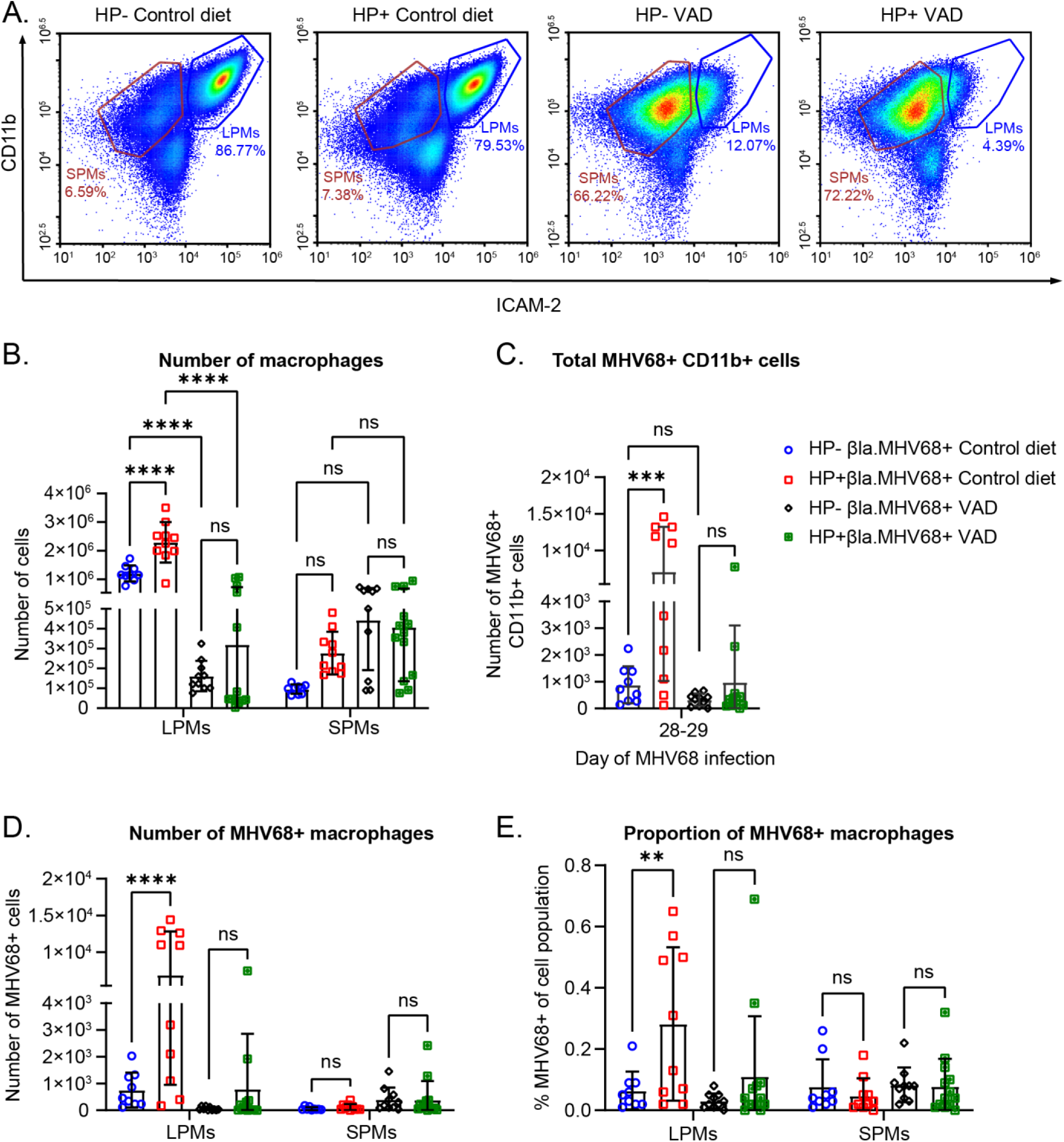
HP-induced increased latent MHV68 infection is dependent on vitamin. **A**. Mice were raised on a vitamin A deficient diet or a control diet, and infected with HP followed by the MHV68.ORF73β-lactamase reporter virus, as in Figure 2. PECs were collected on day 28 or 29 of MHV68 infection. CD11b+ cells were isolated by MACs columns before staining. (A) Representative flow plots of the macrophage populations with the control diet or VAD at day 28 of MHV68 infection. Flow plots are from the single cell gate. (B) Quantification of flow cytometric analysis of LPMs and SPMs at day 28 of MHV68 infection. Data are pooled from 2 independent experiments (n=9-13/group, mean ± standard deviation). Each dot represents an individual mouse. P-values, 2-way ANOVA, Tukey’s multiple comparisons. (C-E) Quantification of flow cytometric analysis of MHV68-infected CD11b+ PECs at day 28 or 29 of MHV68 infection. Data are pooled from 2 independent experiments (n=9-13/group, mean ± standard deviation). Each dot represents an individual mouse. (C) Total number of MHV68-infected CD11b+ PECs. (D) Number of MHV68-infected macrophages. (E) Proportion of MHV68-infected cells out of the parent populations. (D-F) P-values, 2-way ANOVA, Tukey’s multiple comparisons. * P ≤ 0.05, ** P ≤ 0.01, *** P ≤ 0.001, **** P ≤ 0.0001.

### Increased MHV68 infection during HP coinfection requires GATA6-mediated LPM expansion

Although the above experiments suggest that expansion of LPMs were required for increased MHV68 infection, the removal of vitamin A broadly impacts many cells in the immune system. To more specifically target LPMs, we examined mice with macrophage-specific deletion of the transcription factor, GATA6 (*Gata6*^*tm2*.*1Sad*^; *Csf1r*^*MeriCreMer*^). GATA6 is a regulator of large peritoneal macrophages and is required for the transcriptional phenotype and proliferative renewal of LPMs (*20, 29*). GATA6 expression in LPMs is induced by retinoic acid and requires dietary vitamin A. Therefore, mice with macrophage-specific *Gata6* deficiency (*Gata6*^*ΔMac*^) were infected as in Fig. 2A. Peritoneal cells were profiled from CD115-cre+ (*Gata6*^*ΔMac*^) and cre-(*Gata6*^*flox/flox*^) mice at day 2 after MHV68 infection. We observed reduced expression of F4/80 in the ICAM-2+ LPMs in *Gata6*^ΔMac^ mice, similar to previous reports (*9, 20, 29*) (Fig. 5A). We also noted an expansion of the proportion of SPMs in the F4/80+ cells in Gata6^ΔMac^ mice. We therefore gated on the F4/80^hi^ and the F4/80^lo^ ICAM-2+ cells, as well as the SPMs. We detected fewer F4/80^hi^ LPMs in Gata6^ΔMac^ mice compared with littermate controls in MHV68-single infected mice (Fig. 5A and B). We also noted that in contrast to Gata6^flox/flox^ control mice that were coinfected with HP and MHV68, the *Gata6*^*ΔMac*^ mice showed significantly less expansion of the F4/80^hi^ICAM-2^hi^ LPMs, suggesting that HP infection does not efficiently expand LPMs in Gata6^ΔMac^ mice. Although the proportion of SPMs is higher in MHV68-single and HP/MHV68 coinfected Gata6^ΔMac^ mice, the total number of SPMs in all groups was equivalent (Fig. 5B).

**Figure 5.**
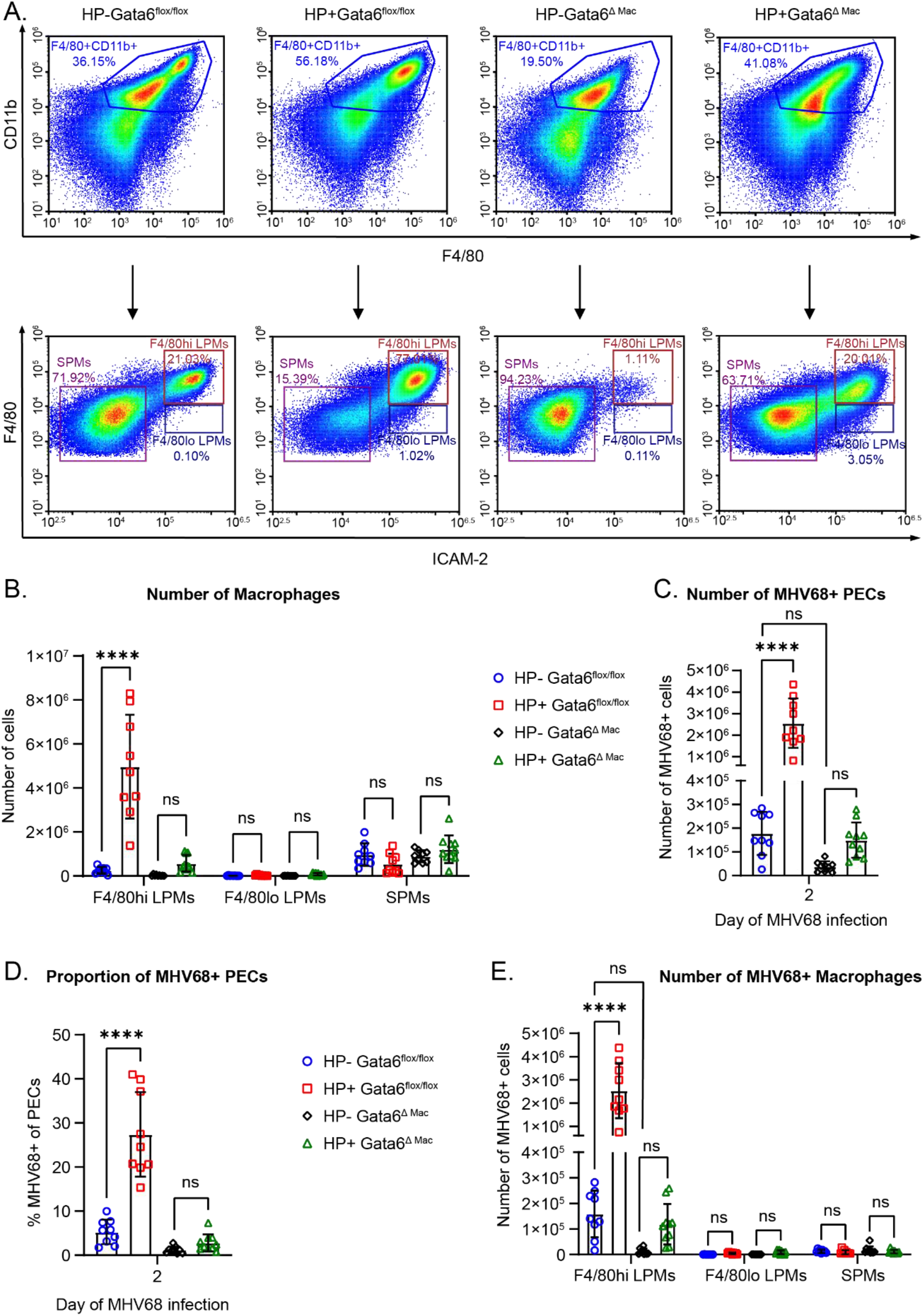
Increased MHV68 infection during HP coinfection requires GATA6-mediated LPM expansion. (A-E) Tamoxifen treated *Gata6*^*flox/flox*^ and *Gata6*^*Δ Mac*^ mice were infected with the MHV68.ORF73β-lactamase reporter virus, as in Figure 2. PECs were collected at day 2 of MHV68 infection for flow analysis. (A) Representative flow plots of macrophage gating at day 2 of MHV68 infection. (B-E) Quantification of flow cytometric analysis of MHV68-infected PECs at day 2 of MHV68 infection. Data are pooled from 2 independent experiments (n=9-10/group, mean ± standard deviation). Each dot represents an individual mouse. (B) Number of macrophages. (C) Total number of MHV68-infected PECs. (D) Proportion of total MHV68-infected PECs. (E) Number of MHV68-infected macrophages. P-values, 2-way ANOVA, Tukey’s multiple comparisons. * P ≤ 0.05, ** P ≤ 0.01, *** P ≤ 0.001, **** P ≤ 0.0001.

We further examined the number and proportion of LPMs that were infected with MHV68 using the MHV68.ORF73β-lactamase virus. As expected at day 2 of MHV68 infection, control Gata6^flox/flox^ mice infected with both HP and MHV68 had a higher proportion and a higher number of virus-positive cells than mice infected with only MHV68 (Fig. 5C and D). In the coinfected Gata6^ΔMac^ mice, we observed minimal increase in the total number of MHV68-positive cells (Fig. 5C and D). When we looked specifically at the F4/80^hi^ or F4/80^lo^ LPMs in Gata6^ΔMac^ mice we did not see increased numbers of infected cells in the coinfected mice (Fig. 5E). There was minimal contribution of MHV68 infection in SPMs (Fig. 5E). Taken together these data suggest that a retinoic acid-dependent transcription factor essential for LPM-specific gene expression and proliferation was required for parasite-induced LPM proliferation and increased virus infection of LPMs.

### Helminth and viral infections alter expression of retinoic acid metabolizing genes in LPMs

Because our data thus far suggested that vitamin A-dependent LPM expansion and the retinoic acid responsive gene, *Gata6*, are required to increase viral infection with HP-coinfection, we questioned whether retinoic acid metabolism was altered in LPMs during helminth/virus coinfection. Therefore, we sorted specifically LPMs on day 1 of MHV68 infection (day 8 of HP infection) from HP-only, MHV68-only, HP/MHV68 coinfected or uninfected mice for gene expression analysis. We found that expression of *Aldh1a3* was increased in HP-infected and HP/MHV68 coinfected mice, while there was no change in MHV68-only infected mice. Expression of *Aldh1a2* and *Aldh1a1* was similar in all groups (Figure 6A).

**Figure 6.**
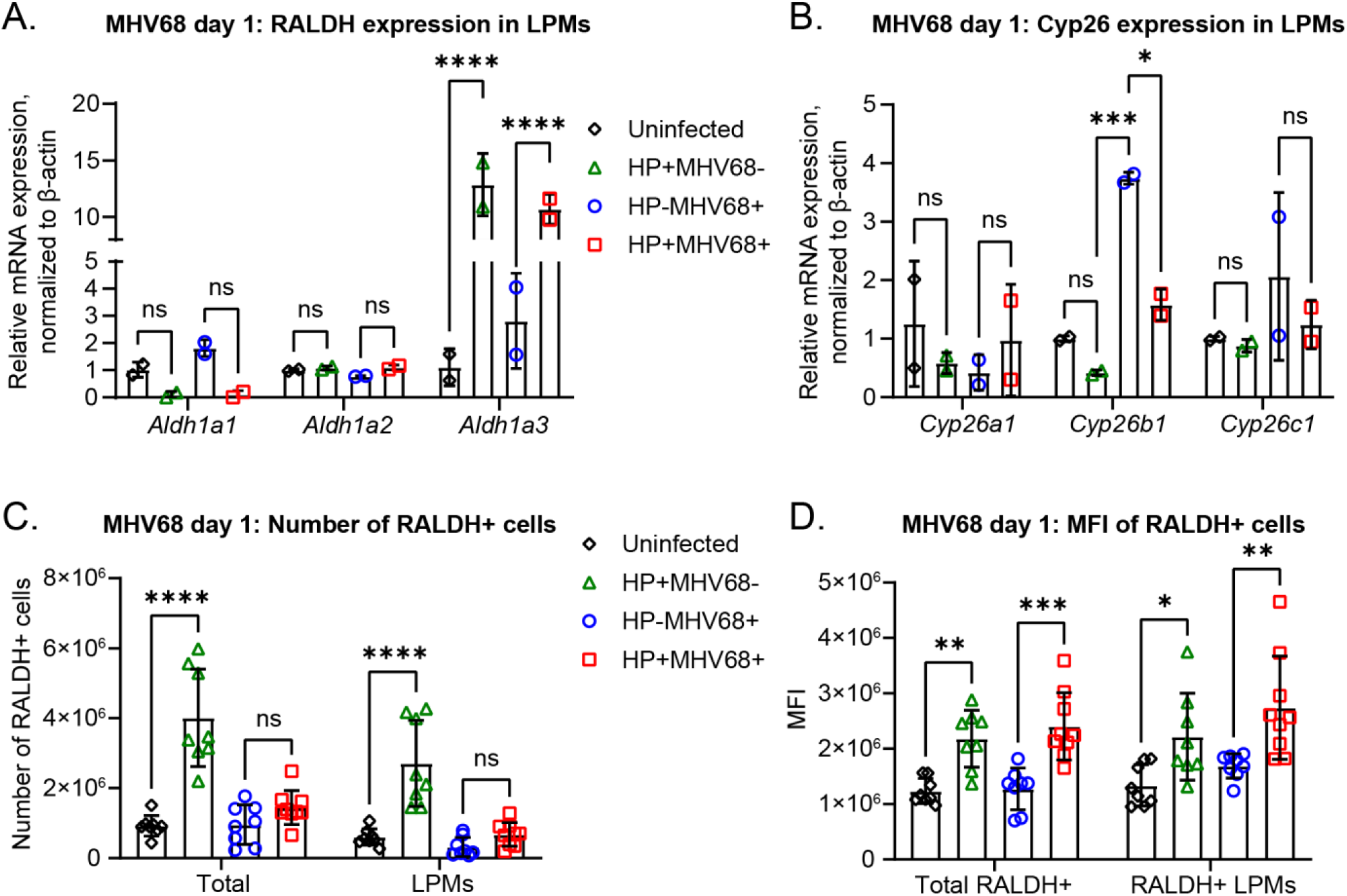
Helminth infection alters expression of retinoic acid metabolizing genes in LPMs. (A-B) HP-infected or uninfected mice were challenged with MHV68 i.p. 7 days later. PECs were collected at day 1 of MHV68 infection. CD11b+ cells were enriched by MACS columns before sorting for LPMs. 4-5 mice/group were pooled for FACS sorting. LPMs were gated as CD3-CD19-Siglec F-CD11b^hi^ ICAM-2^hi^. CT values were normalized to β-actin and then normalized to the delta CT of uninfected LPMs. Data pooled from 2 independent experiments (mean ± standard deviation). (A) Gene expression of retinoic acid producing enzymes in LPMs. (B) Gene expression of retinoic acid catabolizing enzymes in LPMs. (C-D) PECs were collected on day 1 of MHV68 infection and ALDEFLUOR assay was performed. RALDH+ cells were detected based on the ALDEFLUOR fluorescence of the DEAB control sample for each mouse. LPMs were gated as CD3-CD19-Siglec F-CD11b^hi^ ICAM-2^hi^. Data are pooled from 2 independent experiments (8-9 mice/group, mean ± standard deviation). Each dot represents an individual mouse. (C) Number of RALDH+ cells in total PECs and LPMs. (D) Quantification of the Median Fluorescent Intensity of the RALDH+ cells from (C). P-values, 2-way ANOVA, Tukey’s multiple comparisons * P ≤ 0.05, ** P ≤ 0.01, *** P ≤ 0.001, **** P ≤ 0.0001

*Cyp26a* and *Cyp26c* had no significant difference in expression for any of the groups. *Cyp26b1* expression in HP-infected mice was slightly downregulated compared to uninfected mice. In contrast, there was increased expression of *Cyp26b1* in MHV68-only infected mice compared to all the other groups. *Cyp26b1* expression in the coinfected mice was not significantly different from the uninfected group (Figure 6B). Notably, the pattern of *Cyp26* gene expression between uninfected and HP-infected mice was different in these experiments compared with experiments in Figure 1. However, experiments in Figure 1 include all CD115+ PECS, whereas these experiments assessed gene expression in LPMs specifically. Altogether, these data demonstrate that the expression of ATRA metabolizing genes in LPMs was altered during single infection and coinfection states.

The gene expression changes suggest altered production and metabolism of ATRA, which led us to question whether viral infection altered the capacity of PECs to make retinoic acid. We utilized the ALDEFLUOR assay to identify cells with increased capacity for retinoic acid production. In this assay a fluorescent substrate, BODIPY-aminoacetaldehyde, is added to the PECs. This substrate is converted by aldehyde dehydrogenase (RALDH) to BODIPY-aminoacetate, which is unable to diffuse across the cell membrane. RALDH enzymes are responsible for conversion of retinol to retinoic acid, so detection of the fluorescent product by flow cytometry identifies cells that have the capacity to produce retinoic acid.

We analyzed PECs from day 1 of MHV68 infection with the ALDEFLUOR assay. We found that there was an increased total number of RALDH positive (RALDH+) PECs and specifically LPMs from HP-infected mice compared to uninfected mice. We noted a similar increase in RALDH+ PECs and LPMs in HP/MHV68 coinfected mice, although it was not statistically significant (Figure 6C). MHV68-only infected PECs or LPMs showed no change in RALDH staining (Figure 6C). Since we saw a trend for increased RALDH+ cells in both HP-infected an HP/MHV68 coinfected mice, we questioned if the RALDH+ cells had increased enzymatic capacity. To address this, we calculated the median fluorescent intensity (MFI) of the RALDH+ cells. We found that there was increased MFI in the total RALDH+ cells and RALDH+ LPMs of both the helminth-only infected mice and coinfected mice (Figure 6D). This suggests that the RALDH+ cells in HP infected mice produced more retinoic acid than the RALDH+ cells of uninfected or virus-only infected mice. Taken with the gene expression data, we conclude that LPMs from HP-infected and HP/MHV68 coinfected mice have altered ATRA-metabolizing gene expression and increased retinoic acid production capacity.

### Coinfection with *H. polygyrus* increases *ex vivo* MHV68 reactivation in a vitamin A dependent manner

We next determined if the increased infection of MHV68 during helminth coinfection resulted in impaired control of chronic viral infection. Although gammaherpesviruses cause asymptomatic infection in immunocompetent hosts, a key measure of host control of chronic viral infection and pathogenesis is the frequency of viral reactivation from latency (*30*). Notably, cancers driven by human gammaherpesviruses such as Epstein Barr Virus (EBV) and Kaposi’s sarcoma-associated herpesvirus (KSHV) are associated with herpesvirus reactivations from latency (*15*). We therefore examined virus reactivation by limiting dilution assays (LDA) four weeks after MHV68 infection in virus-only and HP/MHV68 coinfected mice (*28*). LDAs quantify the frequency of MHV68 reactivation within a proportion of cells. PECs were plated in serial two-fold dilutions on mouse embryonic fibroblasts and cytopathic effect (CPE) was quantified three weeks later. We found that coinfected mice had a 16-fold higher reactivation frequency in PECs than mice that received only virus (1 out of 1,267 cells versus 1 out of 20,231 cells, respectively) (Figure 7A). We also performed LDA’s with disrupted samples of PECs to quantify any preformed virus that replicated *in vivo*. There was no preformed virus from either group, suggesting that the virus was latent *in vivo* prior to explant (Supplementary Figure 4A).

**Figure 7.**
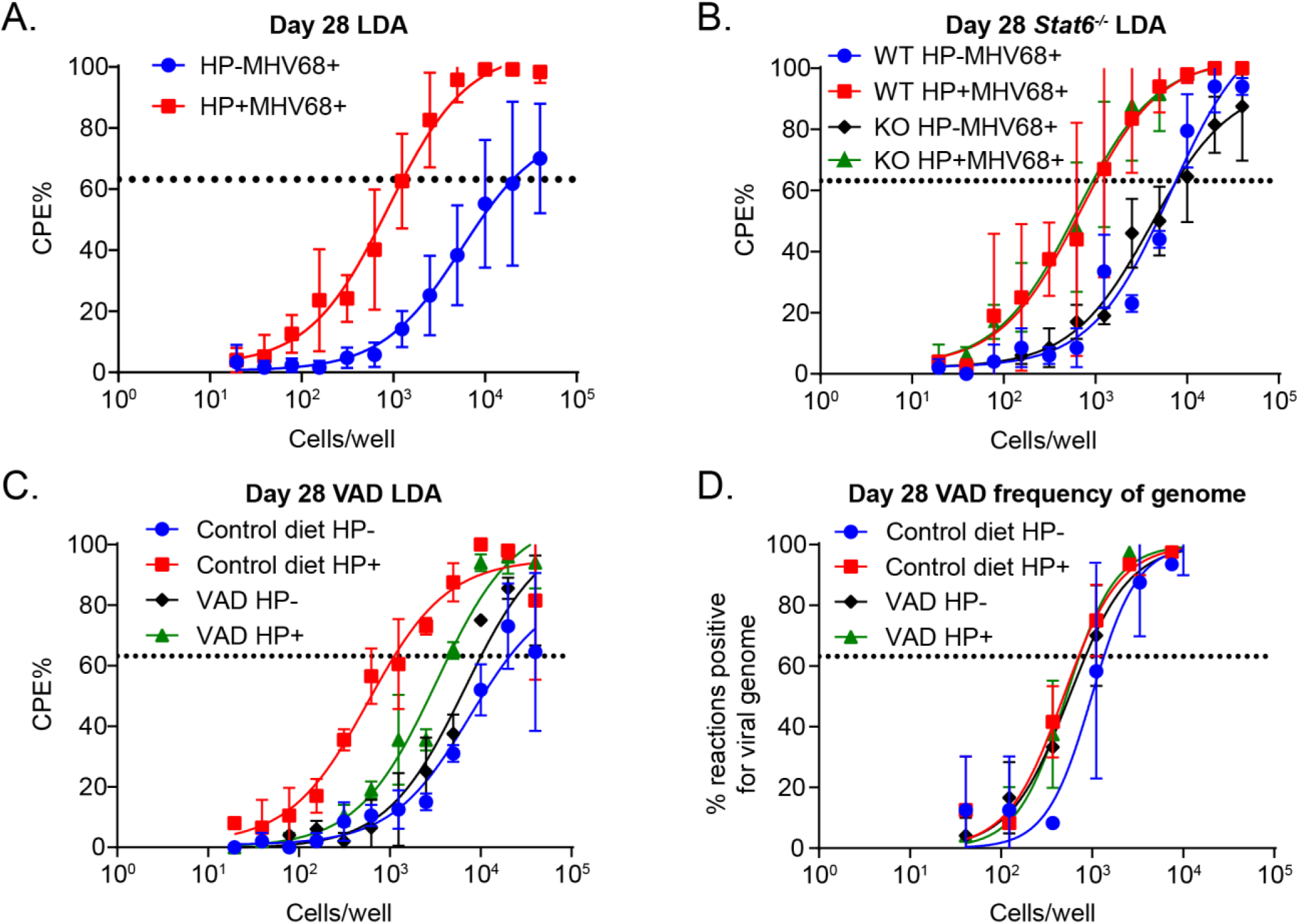
Coinfection with *H. polygyrus* increases *ex vivo* reactivation in a vitamin A-dependent manner. HP-infected or uninfected mice were challenged with 10^6^ PFU of MHV68 i.p. PECs were isolated at day 28-31 of MHV68 infection. (A) Limiting dilution assays were performed from C57BL/6 PECs. Data pooled from 4 independent experiments (3 mice pooled/group). (B) *Stat6*^*-/-*^ or littermate control mice were infected and LDAs were performed at day 28-29 of MHV68 infection. Data are pooled from 2 independent experiments (3 mice pooled/group). (C-D) Mice on VAD or control diet were infected and PECs collected on day 28 of MHV68 infection. LDA’s and LD-PCR were performed on the PECs to determine *ex vivo* reactivation (C) and frequency of viral genomes (D). Data are pooled from 2 independent experiments (3-5 mice pooled/group). Dotted line represents Poisson distribution.

We examined whether increased virus reactivation in mice with helminth infection required STAT6 signaling. We previously showed that challenge with HP during MHV68 latency increased *in vivo* reactivation in a STAT6-dependent manner (*13*). To determine whether this same mechanism was occurring in the coinfected mice that received helminth prior to MHV68 infection, we infected *Stat6*^-/-^ mice and their littermate controls and performed LDAs. We found that coinfected *Stat6*^-/-^ mice still had increased reactivation compared to MHV68-infected *Stat6*^-/-^ mice (Figure 7B). The *Stat6*^-/-^ mice also did not have any preformed virus in disrupted samples (Supplementary Figure 4B). This confirmed that STAT6 signaling was not driving the increased reactivation in coinfected mice when HP infection occurs before MHV68 infection.

Since we demonstrated that the helminth-induced increase in infection was dependent on vitamin A, we questioned whether that was true for the helminth-induced reactivation. Coinfected mice on the control diet had an 18-fold increase in reactivation compared to the virus-only control diet mice (1 out of 20,930 in MHV68-only versus 1 out of 1124 in coinfected). In contrast, coinfected mice on the VAD diet had only a 2-fold helminth-induced increase in reactivation compared to virus only mice on the VAD diet (1 out of 4,479 in coinfected VAD versus 1 out of 10,089 in virus-only VAD) (Figure 7C). Disruption of samples confirmed that the virus was latent and there was no *in vivo* reactivation in any of the groups (Supplementary Figure 4C). To ensure that loss of vitamin A did not disrupt the establishment of latency, we performed LD-PCR to detect viral genome and confirmed that latency establishment was equivalent in the PECs for all groups (Figure 7D). Overall, these results demonstrate that coinfection with a helminth induces increased *ex vivo* reactivation of MHV68 in a vitamin A dependent manner.

## Discussion

The purpose of this study was to determine the impact of intestinal parasite infection on tissue resident macrophages in the peritoneal cavity and their response to viral infection. We found that HP parasite infection expands LPMs. This proliferation of LPMs did not require STAT6 signaling but did require retinoic acid. We further determined that parasite coinfected mice have increased numbers of herpesvirus infected LPMs, during both acute replication and latency, and this expanded infection required dietary vitamin A and the transcription factor GATA6. We also showed that LPMs from both HP-only and HP/MHV68 coinfected mice have altered expression and activity of retinoic acid metabolizing genes. Last, we demonstrated that coinfected mice have increased *ex vivo* reactivation of MHV68, which was dependent on vitamin A. These results suggest that intestinal parasite infection expands tissue resident macrophages in the peritoneal cavity and that this leads to increased viral infection and herpesvirus reactivation from these cells (Figure 8).

**Figure 8.**
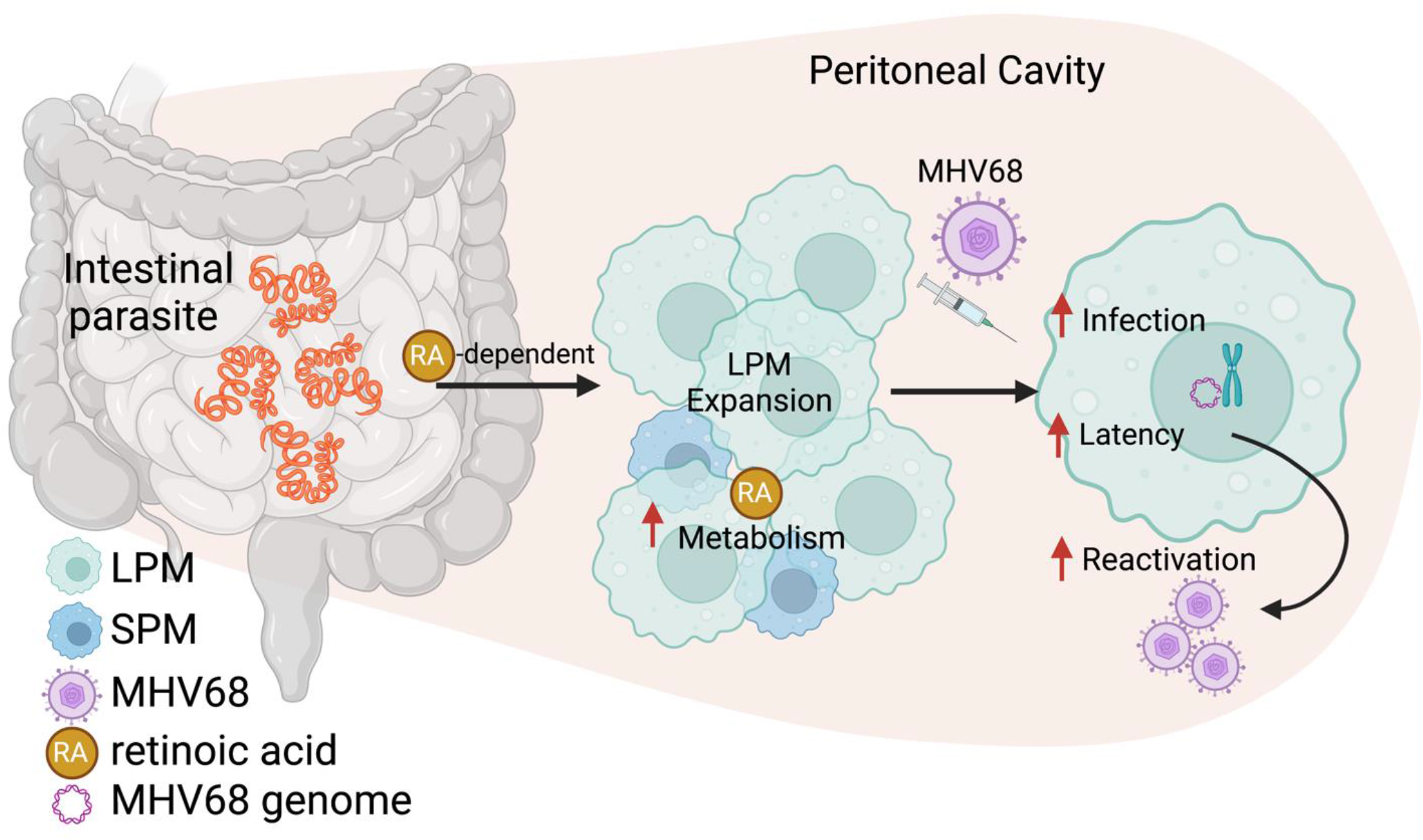
Model: Intestinal helminth infection induces LPM expansion in a retinoic acid dependent manner. Expanded LPMs leads to increased infection and reactivation of MHV68. Created in BioRender.

We demonstrated that LPMs expanded during early timepoints in HP infection, and that this expansion required retinoic acid. While other studies have demonstrated that IL-4 receptor signaling is sufficient to induce LPM or serous cavity macrophage expansion (*7, 8, 19*), we found that STAT6, the transcription factor downstream of the IL-4 receptor, was not necessary for HP-induced LPM expansion. This indicated that HP infection induced LPM expansion through other signals than IL-4 or IL-13 in our system. Such signals could be immune derived, as has been described before (*7, 8, 19*). Because retinoic acid is important for the maintenance and proliferation of LPMs (*20*), we tested whether blockade of retinoic acid-induced gene expression inhibited LPM expansion after HP infection. Our data revealed that retinoic acid was indeed important for parasite-induced LPM expansion. Another option is that there is also a signal for LPM expansion that comes directly from HP. HP has been shown to secrete immunomodulatory molecules (*1*–*3*) which leaves the possibility that there could also be an HP secreted signal that induces LPM expansion.

Our results reported here compared with our previous study on HP/MHV68 coinfection reveal that the timing of coinfection dictates mechanism. We previously discovered that when mice with a latent MHV68 infection were challenged with HP four to six weeks after the initial viral infection, the HP infection induced reactivation *in vivo* (*13*). This reactivation required STAT6 signaling, suggesting that IL-4/IL-13 produced in response to the HP infection promoted virus reactivation. In contrast, in this study we first infected mice with HP and challenged with MHV68 seven days later. While there is still increased reactivation of the virus in HP-infected mice, the mechanism did not require STAT6 signaling. Together, these two systems reveal that the order and timing of coinfection are critical variables in the determination of mechanism.

Coinfection is a multidimensional problem, where each infection potentially impacts the other infection. We primarily focused on the impact of helminth infection on the macrophage response in the peritoneal cavity and on the replication and latency of MHV68. However, it is unknown if MHV68 infection impacts the helminth infection. We measured the numbers of adult worms and eggs produced in the stool of HP-single infected and HP/MHV68 coinfected mice and found no differences in worm burden or fecundity. This does not eliminate the possibility that there could still be differences in the host response to helminth infection in the intestine. In addition, we have not assessed the impact of LPM expansion specifically on intestinal helminth infection. LPMs are important for IgA production by B1 B cells and secretion of IgA in the feces (*20*). Because antibodies are required for protective immunity against secondary HP infections (*31*), an interesting question for future studies is whether expanded LPMs promote antibody responses against HP. Large cavity macrophages may also play a direct role in controlling worm infection. Increased numbers of large cavity macrophages in the pleural cavity are correlated with reduced burden of *L. sigmodontis*, a filarial lung nematode(*11*).

An intriguing observation is that helminth infection appears to buffer the “macrophage disappearance reaction (MDR)”. Although our data agrees with other reports that parasites expand LPMs, other types of infections, such as bacterial infection, trigger MDR in the peritoneal cavity (*32*). The MDR causes LPMs to migrate to the omentum (*20*), the periphery of the peritoneal cavity, or to form clots (*32*). Viral infection in the peritoneal cavity also causes MDR, because analysis of the peritoneal cell populations during acute replication of MHV68 showed that LPMs were depleted by day 8 of infection and reappeared by day 14 (*16*). This is consistent with our observations, where MHV68-only infected mice lose LPMs during infection but have a prominent LPM population during latency. Perhaps more intriguing, is the fact that coinfected mice did not lose their LPM population, which suggests that the helminth infection helps maintain the LPM population during acute inflammation in the peritoneal cavity. Humans, unless they currently live in developed countries, are infected with parasites throughout their lives. We question whether tissue resident macrophage populations are maintained in individuals with intestinal parasite infections, which may confer susceptibility or resistance to infections of other tissues.

Finally, our data indicate that early alterations in macrophage phenotype and numbers have dramatic effects on herpesvirus latency and reactivation. Although we saw increased viral infection of LPMs, we did not detect significant differences in acute replication by plaque assay. Our data supports the hypothesis that some herpesvirus genomes that enter cells immediately become latent, rather than replicating, as was seen previously using the MHV68.ORF73β-lactamase virus (*16*). We hypothesize that MHV68 may be sensitive to altered retinoic acid levels in LPMs at early infection timepoints. Our expression data from the sorted LPMs suggests that RA metabolism may be altered by intestinal helminth infection. The local environment of tissue resident macrophages influences the enhancer landscape of the macrophages, and LPMs cultured with RA maintain some of the enhancers on LPM-specific genes (*33*). If RA impacts the epigenetic markings of the host cell, it is possible that RA may induce alterations of the epigenetic markings of the MHV68 episome, which resides in the nucleus during latency (*34*). In fact, RA was shown to induce MHV68 gene expression in bone marrow derived macrophages (*12*), implicating RA in the regulation of MHV68 latency and reactivation.

In summary, we demonstrate that HP infection promoted LPM expansion in a retinoic acid-dependent manner. When we challenged parasite-infected animals with a systemic herpesvirus infection, we discovered that HP coinfected animals maintain the LPM population during the secondary MHV68 challenge, which normally causes MDR. We found that the expansion of LPMs by helminth infection was required for increased herpesvirus reactivation from latency, because coinfected mice depleted of LPMs by vitamin A dietary deficiency did not have increased virus reactivation. Taken together our results indicate that intestinal parasite infection alters tissue resident macrophages in non-intestinal tissues, which can increase the cellular reservoir for herpesvirus infection. By expanding the cellular reservoir, the parasite coinfection increases herpesvirus reactivation from latency.

## Author contributions

CMZ, CD, and TAR designed experiments. CMZ performed experiments and analyzed data. CMZ and TAR wrote and edited the manuscript. CD, MP, and LVH gave critical scientific feedback and shared reagents and mice. JC assisted with experiments. PD assisted with helminth propagation. TAR secured funding.

## Acknowledgements

We thank the UTSW Animal Resource Center, UTSW IACUC, UTSW Flow Core, and Moody Foundation Flow Cytometry Core for use of their facilities. We thank Jaime Vaquer-Alicea for his assistance with flow sorting. We thank Dr. Kun Yang, Dr. Conor Finlay, and James Parkinson for their tamoxifen protocols. We thank Dr. Scott Tibbetts for MHV68.ORF73βla. We thank Dr. Gwendalyn Randolph for the *Gata6*^f/f^ and CD115 Cre-ER strains and genotyping protocols.

## Methods

### Mice

Animals were bred and housed in pathogen-specific free facilities. All procedures were approved by the UT Southwestern Medical Center IACUC. Mice were bred in a barrier facility and experiments were performed in a conventional facility. Unless noted, C57BL/6J mice were used in the experiments. *Stat6*^-/-^ (B6.129S2(C)-*Stat6*^*tm1Gru*^/J, stock number 005977) (*35*) mice were purchased from Jackson Labs and bred to C57BL/6 mice to generate littermate controls or maintained in a *Stat6*^-/-^ homozygous cross. *Gata6*^f/f^ (*Gata6*^*tm2*.*1Sad*^/J, Jackson stock number 008196) (*36*) and CD115 Cre-ER (Tg(*Csf1r*-cre/Esr1*)1Jwp/J, Jackson stock number 019098) (*32, 37, 38*) mice were a gracious gift from Dr. Gwendalyn Randolph. *Gata6*^f/f^ mice were crossed with CD115 Cre-ER mice, so that each litter produced cre- and cre+ littermate controls. To delete *Gata6* from macrophages and monocytes, tamoxifen (Sigma # T5648) was dissolved at 20 mg/mL in autoclaved corn oil (Sigma # C8267) by shaking overnight at 37 degrees C in the dark. Mice were orally gavaged with 4 mg of tamoxifen (200 µL of 20 mg/mL stock) dissolved in corn oil for three consecutive days. Experiments utilized a mix of both male and female mice. Mice were aged 5-14 weeks old at the start of infections.

### Animal infections

Mice were orally gavaged with 100 L3 *H. polygyrus* one week before infection with MHV68. Mice were infected with 1 × 10^6^ pfu of MHV68 (WUSM strain) intraperitoneally. MHV68.ORF73β-lactamase was a gift from Dr. Scott Tibbetts and has been previously described (*26*).

### *H. polygyrus* (HP) maintenance

C57BL/6 or *Stat6*^-/-^ mice were infected with 200-400 L3 HP by oral gavage. Two weeks after infection, grates were placed in the bottom of the cages and feces were collected for 3-5 days. Feces were plated with washed and autoclaved deactivated charcoal (C2764, Sigma). 7-10 days after fecal plating, L3 larva were collected using a Baermann apparatus as previously described (*39*). Larva were stored at 4°C in PBS for up to 6 months.

### HP worm burden and fecundity

Small intestines were collected at the time of euthanasia. Small intestines were cut open longitudinally and incubated at 37°C for 1-5 hours in PBS. Adult worms were counted by eye. For fecundity, fecal pellets were collected from the colon at the time of euthanasia. The fecal pellets were suspended (or dissolved) in water. An equal volume of saturated sodium chloride (Sigma) was added and eggs were enumerated on a McMaster slide (Electron Microscopy Sciences).

### Vitamin A deficient diet model

Female C57BL/6 breeders were given regular chow until day 14 of gestation, as described (*40*). The females were then given either control diet (Teklad, TD.09838) or vitamin A deficient diet (Teklad, TD.09839) until the pups were weaned. Weaned pups were fed their respective diets throughout the experiment until they were sacrificed. Mice were housed with Sani-chip bedding to avoid any uncontrolled vitamin A intake.

### BMS 493 treatment

BMS 493 (Tocris, Cat #3509) was dissolved at a concentration of 7.33 mg/mL in DMSO. 30 µL of stock BMS 493 was diluted with 70 µL of olive oil, per mouse. Mice were gavaged with 220 µg BMS 493 or the vehicle control (DMSO in olive oil) every other day until euthanasia.

### MHV68 Plaque assays

PECs were harvested by peritoneal lavage with 10 ml of complete D5 ((DMEM (Corning), 5% FBS (Biowest), 2mM L-glutamine (Gibco), 1% HEPES (Corning), 10 U/mL Penicillin and 10 µg/mL streptomycin sulfate (Corning)) and washed once. The PECs were resuspended in 1 ml of D5, then were homogenized with 1 mm zirconia beads (BioSpec Products) in a Precellys 24 (Bertin Instruments) at 5000 rpm for 1 min. 10-fold serial dilutions were plated on 3T12s. Virus was absorbed for 1 hour at 37 degrees C before 1% (w/v) methylcellulose (Sigma-Aldrich) (10 g methylcellulose in 1L MEM (Corning)) was added as an overlay. Plates were incubated at 37 degrees C in 5% CO_2_ for 7 days before fixing with 2% formaldehyde and staining with 0.1% crystal violet (Sigma-Aldrich).

### Limiting dilution assays (LDAs)

Limiting dilution assays were performed as previously described (*28*). Briefly, peritoneal exudate cells (PECs) were collected by lavage in 10 ml of complete D10 (DMEM (Corning), 10% FBS (Biowest), 2mM L-glutamine (Gibco), 1% HEPES (Corning), 10 U/mL Penicillin and 10 µg/mL streptomycin sulfate (Corning)) and washed once. Cells were counted. 2 × 10^6^ PECs were used for the 0 dilution. Two-fold dilutions were performed and plated on C57BL/6 mouse embryonic fibroblasts. For disrupted samples, 2 × 10^6^ PECs were homogenized in a Precellys 24 (Bertin Instruments) at 6000 rpm for 1 min. Two-fold dilutions were plated of the disrupted samples. The assay was scored for cytopathic effect three weeks after plating.

### Limiting dilution (LD)-PCR

Samples from the LDAs were frozen and utilized for LD-PCR. LD-PCR was performed as previously described (*24*). Briefly, cells were counted and 8 × 10^5^ cells (including dead cells) were used to make 3-fold dilutions. The dilutions were used in nested-PCR reactions for ORF72 of MHV68. The proportion of positive reactions was used to quantify the frequency of latency. For samples that were sorted before LD-PCR, 3 × 10^5^ cells starting cells were used. Primers for ORF72:

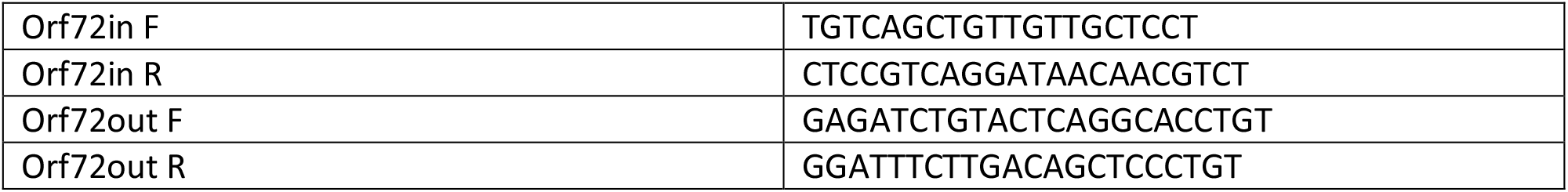

### Flow cytometry

Peritoneal exudate cells (PECs) were collected by lavage with 10 mL of flow buffer (1% FBS in PBS (Corning)). PECs were washed once, then Fc receptors were blocked with α-CD16/32 (Biolegend). The following antibodies were used in various experiments: Alexafluor 647-α-CD102 (3C4 (MIC2/4), Biolegend), APC-Cy7-α-CD19 (1D3, BD Biosciences), redFluor 710-α-CD11b (M1/70, Tonbo), Fitc-α-CD11b (M1/70, BD Biosciences), PE-Cy7-α-CD19 (1D3, Tonbo), Violetfluor450-α-CD3 (17A2, Tonbo), and BV711-α-Siglec F (E50-2440, BD Biosciences). Cells were fixed with 2% formaldehyde (VWR). Samples were run on a Novoctye 3000 (ACEA Biosciences) and analyzed with either Flowjo (Tree Star, Inc, San Carols, CA) or NovoExpress (version 1.5.0, Agilent Technologies, Inc.).

### MHV68.β-lactamase detection by flow cytometry

PECs were collected and washed once. Cells were counted and CD11b+ cells were selected by MACS column purification using the CD11b MicroBead kit and LS columns (Miltenyi Biotec). CD11b+ cells were Fc blocked with α-CD16/32 (Biolegend) and stained with Alexafluor 647-α-CD102 (3C4 (MIC2/4), Biolegend), APC-Cy7-α-CD19 (1D3, BD Biosciences), redFluor 710-α-CD11b (M1/70, Tonbo) or APC-Cy7-α-CD11b (M1/70, Biolegend), and R718-α-CD102 (3C4 (MIC2/4), BD Biosciences). Cells were counted again and resuspended in 3 × 10^6^ cells/mL before adding CC4F-AM (LiveBLAzer FRET-B/G Loading Kit with CCF4-AM, ThermoFisher Scientific). Cells were incubated for 1 hour, then washed with PBS and fixed with 2% formaldehyde. Samples were immediately run on a Novocyte 3000 (ACEA Biosciences).

### ALDEFLUOR assay

PECs were collected, washed once, and then counted. 1 million cells were resuspended in ALDEFLUOR assay buffer, and then the ALDEFLUOR kit (Stemcell Technologies) protocol was followed, with the exception that 10 mM DEAB (Sigma), made in 200 proof ethanol (Sigma), was used instead of the kit’s DEAB reagent. PECs were incubated for 45 min at 37 degrees C, then washed and stained with α-CD16/32 (Biolegend), PE-Cy7-α-CD11b (M1/70, Tonbo), Alexafluor 647-α-CD102 (3C4 (MIC2/4), BD Biosciences), BV570-α-CD19 (6D5, Biolegend), BV711-α-Siglec F (E50-2440, BD Biosciences), APC-Cy7-α-CD45 (30-F11, BD Biosciences), and Violetfluor 450-α-CD3 (17A2, Tonbo). Cells were immediately run on a Novocyte 3000 (ACEA Biosciences).

### Flow sorting

Three to four mice were pooled for each sample. PECs were collected and washed. Cells were counted and CD11b+ cells were selected by MACS column purification using the CD11b MicroBead kit and LS columns (Miltenyi Biotec). CD11b+ cells were stained with Fc block α-CD16/32 (Biolegend) and then PE-Cy7-α-CD19 (1D3, Tonbo), Fitc-α-CD11b (M1/70, BD Biosciences), BV510-α-F4/80 (BM8, Biolegend), BV711-α-Siglec F (E50-2440, BD Biosciences), Violetfluor 450-α-Ly-6G (GR1) (RB6-8B5, Tonbo), and Alexafluor 647-α-CD102 (3C4 (MIC2/4), Biolegend). Samples were run on a BD FACS Aria II SORP and collected in either D10 media (for LD-PCR) or flow buffer (for qPCR). Afterward, the samples were counted and frozen for LD-PCR (in media) or RNA extraction (RNAqueous Lysis buffer).

### qPCR

Either CD115+ PECs were isolated with LS MACs columns (Miltenyi) or PECs were sorted for LPMs before RNA extraction. RNA extraction from sorted LPMs was performed using a RNAqueous-Micro Total RNA Isolation Kit (Invitrogen) according to the protocol. RNA was reverse transcribed using Superscript Vilo cDNA Synthesis kit (Invitrogen). qPCR was performed using PowerUp SYBR Green Master Mix (Applied Biosystems) and cycled on a QuantStudio 7 Flex Real-Time PCR System (Applied Biosystems). Primers for *Cyp26a1, Cyp26b1*, and *Cyp26c1* were taken from Takeuchi, et al. (*41*) and are listed below. Primers for *Arg1* were taken from Okabe, et al. (*20*) and are listed below. Primers for *Aldh1a1, Aldh1a2*, and *Aldh1a3* were designed with the PrimerQuest Tool (IDT). Primers for Beta-actin were designed by the Hooper lab.

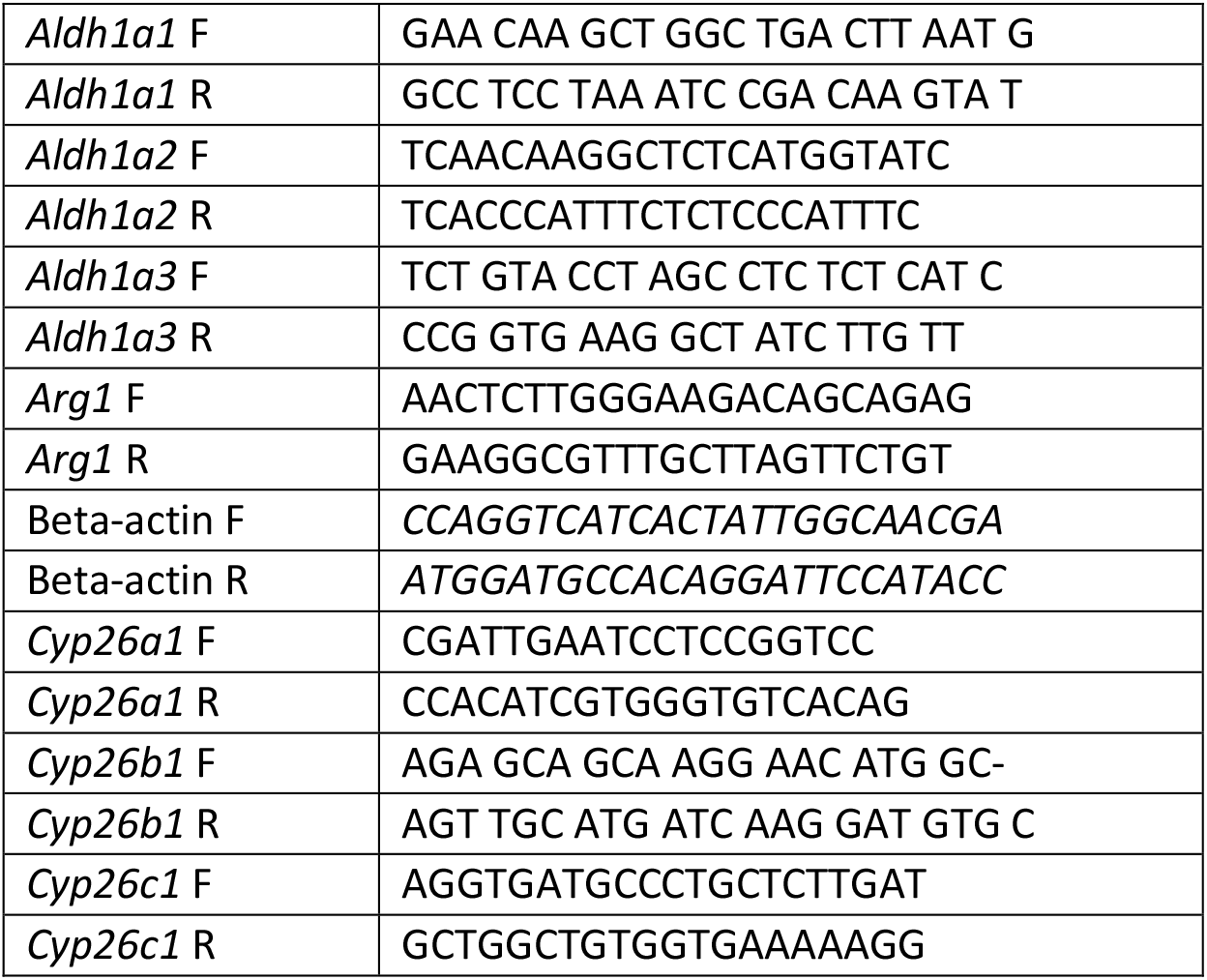

### Statistics

Analysis was performed with GraphPad Prism 9.3.1. An unpaired t-test was used to analyze data with wo groups. One-way or 2-way ANOVA was used to analyze data with more than two groups, with Tukey’s multiple comparisons. Linear regression analysis was used to create the curves for the LDA and LD-PCR graphs. Frequencies of reactivation or latent genome were calculated using the linear regression equations provided by GraphPad Prism. Error bars depict the standard deviation. Significance was considered significant when *P* ≤ 0.05.

## Supplemental Figures

**Supplementary Figure 1.**
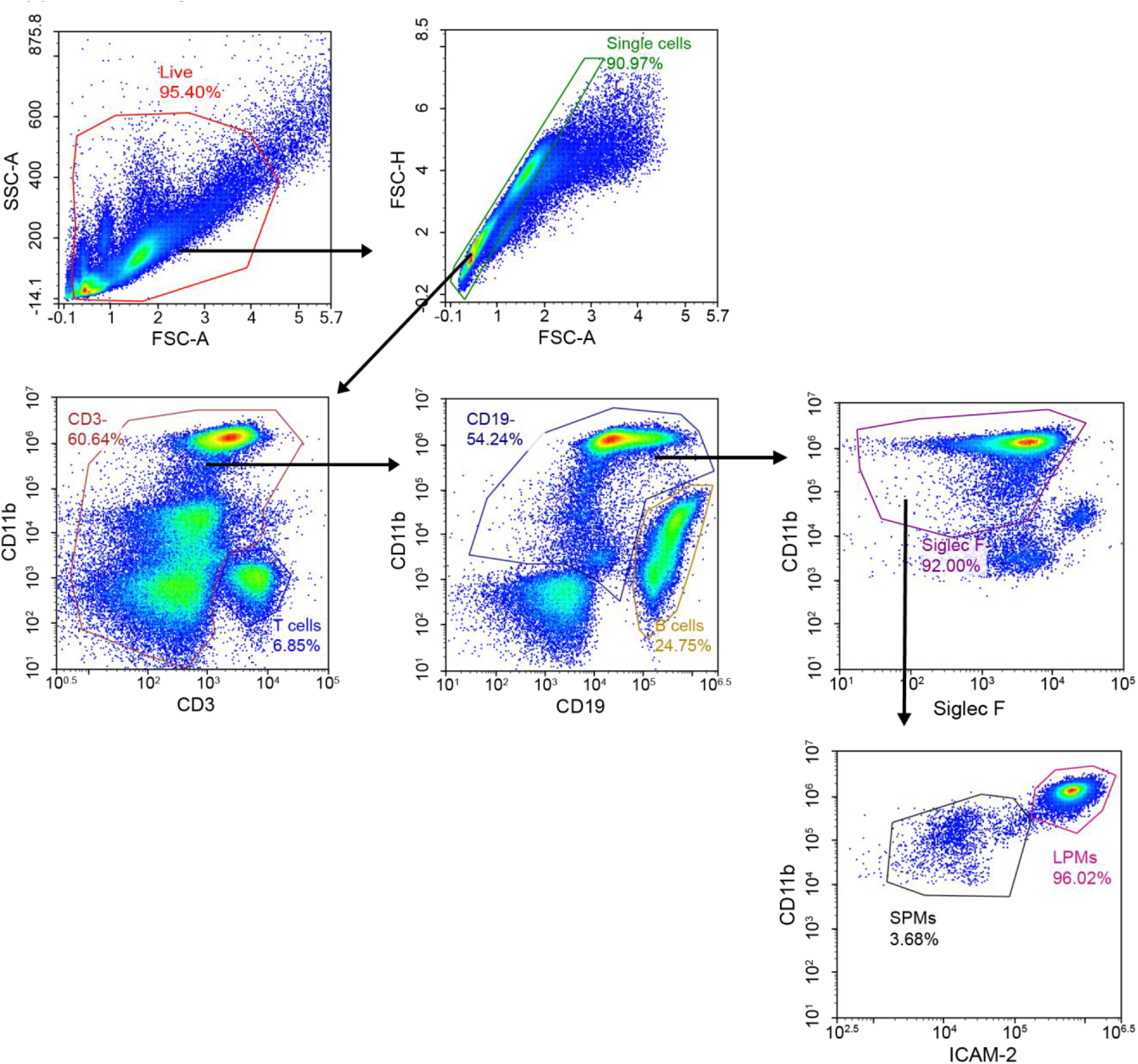
Gating strategy for LPMs.

**Supplementary Figure 2.**
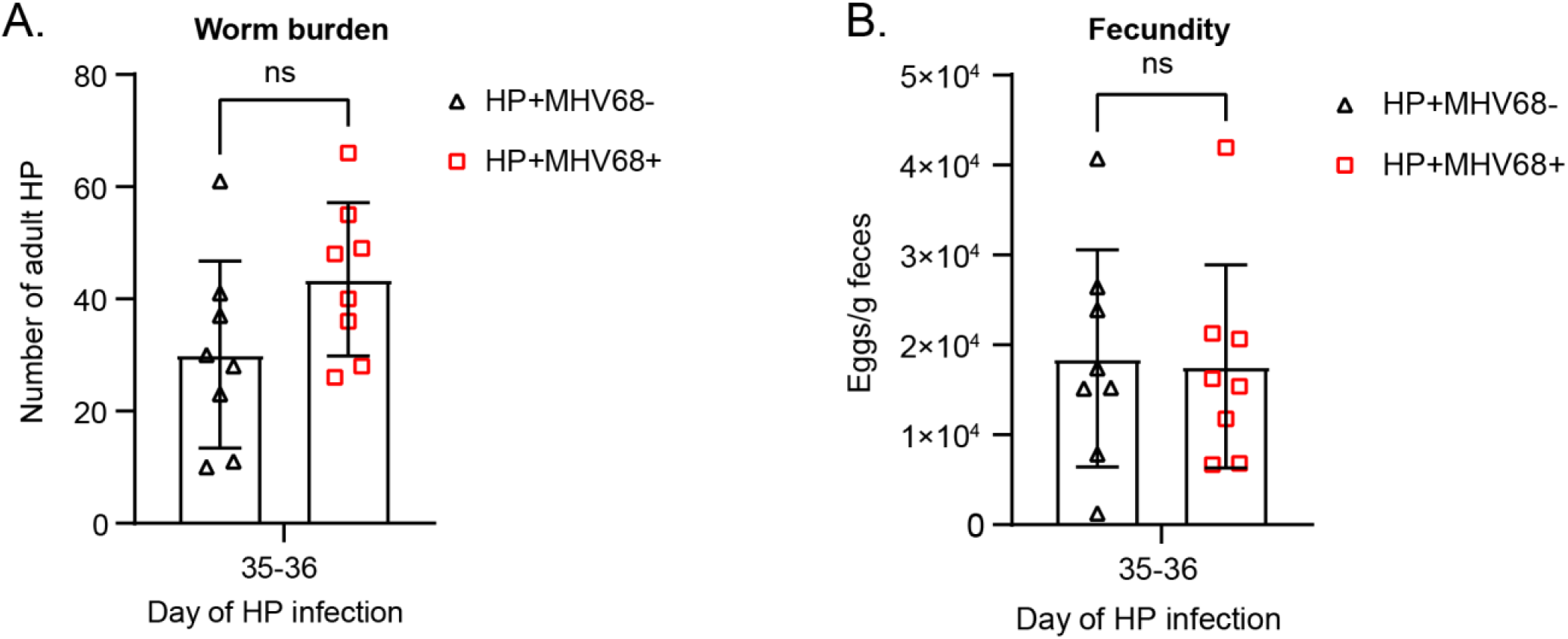
Impact of MHV68 on HP infection. (A-B) Worm burden and fecundity of HP in helminth-only and coinfected mice at days 35 and 36 of HP infection, which correspond to days 28 and 29 of MHV68 infection. Data are pooled from 2 independent experiments (n=8 mice/group, mean± standard deviation). Each dot represents an individual mouse. (A) Adult HP worm burden. (B) Fecundity of HP. P-values, Unpaired t-test. * P ≤ 0.05, ** P ≤ 0.01, *** P ≤ 0.001, **** P ≤ 0.0001.

**Supplementary Figure 3.**
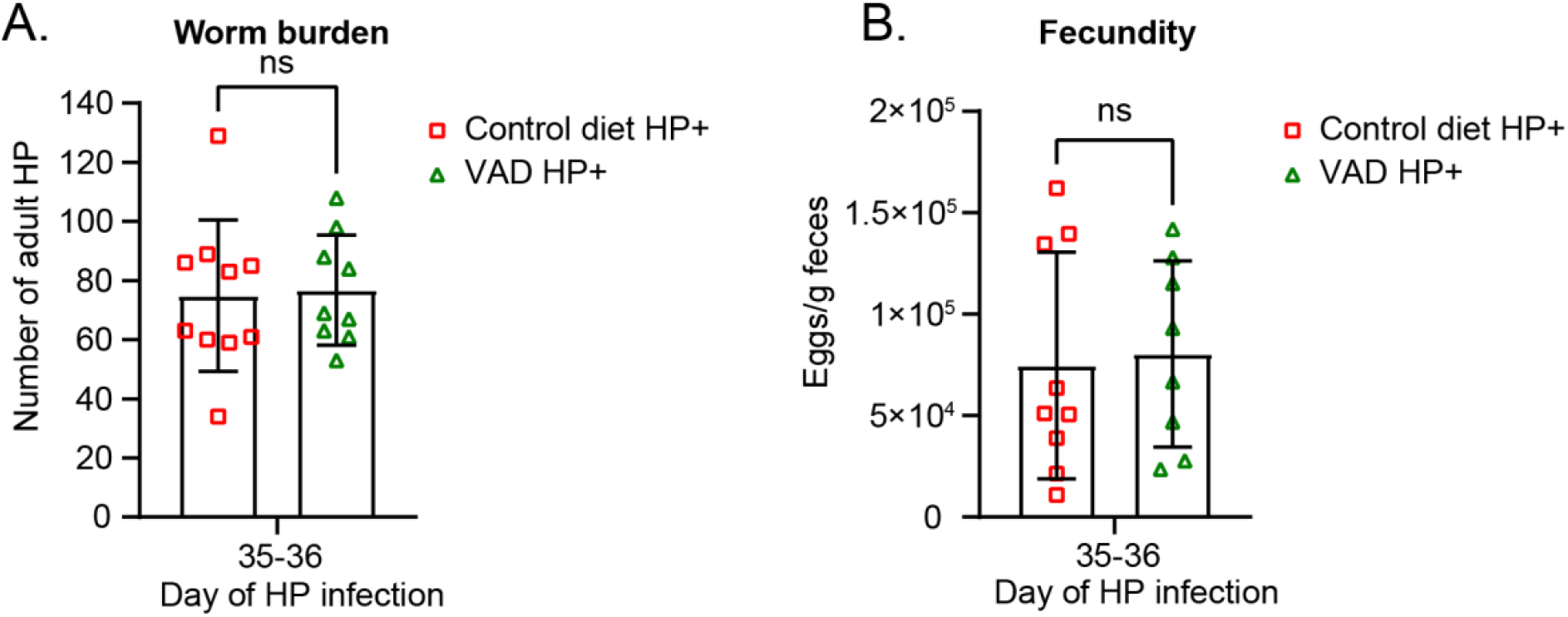
Effect of VAD on HP infection. (A-B) Mice were raised on a vitamin A deficient diet or a control diet, and infected with HP and the MHV68.ORF73β-lactamase reporter virus, as in Figure 2. Worm burden and fecundity of HP at days 35 and 36 of HP infection, which correspond to days 28 and 29 of MHV68 infection. Data are pooled from 2 independent experiments (4-6 mice/group, mean± standard deviation). Each dot represents an individual mouse. (A) Adult HP worm burden. (B) Fecundity of HP. P-values, unpaired t-test. * P ≤ 0.05, ** P ≤ 0.01, *** P ≤ 0.001, **** P ≤ 0.0001.

**Supplementary Figure 4.**
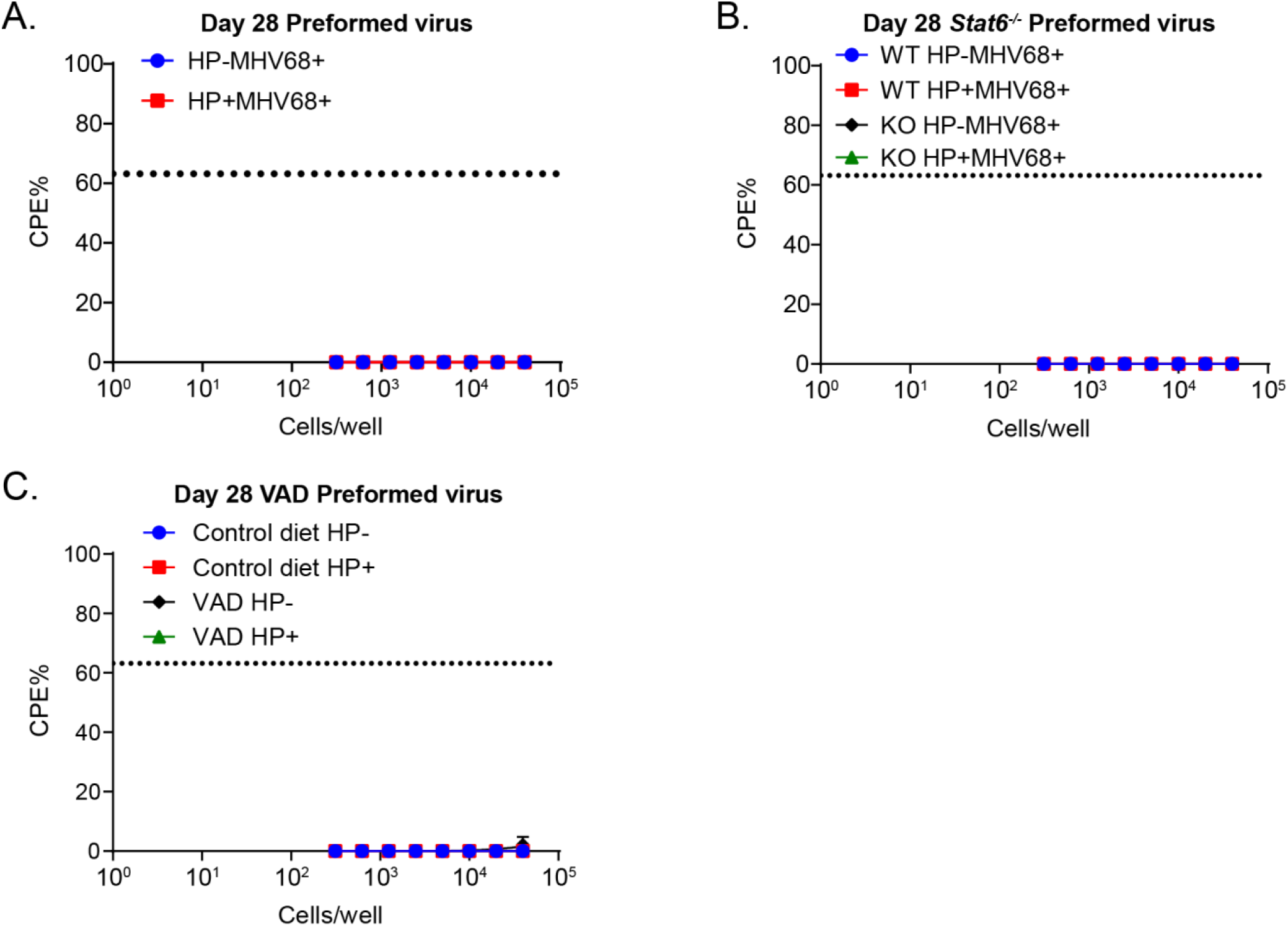
LDAs of preformed virus. HP-infected or uninfected mice were challenged with 10^6^ PFU of MHV68 i.p. PECs were isolated at day 28-31 of MHV68 infection. (A) C57BL/6 PECs were collected at day 28-31 of MHV68 infection for LDAs. PECs from (Figure 7A) were disrupted before plating to detect preformed virus. Data pooled from 4 independent experiments (3 mice pooled/group). (B) Littermate controls of *Stat6*^*-/-*^ PECs from (Figure 7B) were disrupted before plating to detect preformed virus. Data are pooled from 2 independent experiments (3 mice pooled/group). (C) Mice on VAD or control diet were infected as previously described and PECs collected on day 28 of MHV68 infection. PECs from (Figure 7C) were disrupted before plating to detect preformed virus. Data are pooled from 2 independent experiments (3-5 mice pooled/group). Dotted line represents Poisson distribution.

